# An outbreak of SARS-CoV-2 with high mortality in mink (*Neovison vison*) on multiple Utah farms

**DOI:** 10.1101/2021.06.09.447754

**Authors:** Chrissy Eckstrand, Tom Baldwin, Mia K Torchetti, Mary Lea Killian, Kerry A. Rood, Michael Clayton, Jason K. Lott, Rebecca M Wolking, Daniel S Bradway, Timothy Baszler

**Author notes:** Additional author notes should be indicated with symbols (e.g., for equal contributions or current addresses).

## Abstract

The breadth of animal hosts that are susceptible to severe acute respiratory syndrome coronavirus 2 (SARS-CoV-2) and may serve as reservoirs for continued viral transmission are not known entirely. In August 2020, an outbreak of SARS-CoV-2 occurred in multiple mink farms in Utah and was associated with high mink mortality and rapid viral transmission between animals. The outbreak’s epidemiology, pathology, molecular characterization, and tissue distribution of virus within infected mink is provided. Infection of mink was likely by reverse zoonosis. Once established, infection spread rapidly between independently housed animals and farms, and caused severe respiratory disease and death. Clinical signs were most notably sudden death, anorexia, and increased respiratory effort. Gross pathology examination revealed severe pulmonary congestion and edema. Microscopically there was pulmonary edema with moderate vasculitis, perivasculitis, and fibrinous interstitial pneumonia. Reverse transcriptase polymerase chain reaction (RT-PCR) of tissues collected at necropsy demonstrated the presence of SARS-CoV-2 viral RNA in multiple organs including nasal turbinates, lung, tracheobronchial lymph node, epithelial surfaces, and others. Whole genome sequencing from multiple mink was consistent with published SARS-CoV-2 genomes with few polymorphisms. The Utah mink SARS-CoV-2 strain fell into Clade GH, which is unique among mink and other animal strains sequenced to date and did not share other spike RBD mutations Y453F and F486L found in mink. Localization of viral RNA by *in situ* hybridization revealed a more localized infection, particularly of the upper respiratory tract. Mink in the outbreak reported herein had high levels of virus in the upper respiratory tract associated with mink-to-mink transmission in a confined housing environment and were particularly susceptible to disease and death due to SARS-CoV-2 infection.

**Author Summary:** The recent emergence and worldwide spread of the novel coronavirus has resulted in worldwide disease and economic hardship. The virus, known as SARS-CoV-2 is believed to have originated in bats and has spread worldwide through human-to-human virus transmission. It remains unclear which animal species, other than humans, may also be susceptible to viral infection and could naturally transmit the virus to susceptible hosts. In this study, we describe an outbreak of disease and death due to SARS-CoV-2 infection in farmed mink in Utah, United States. The investigation reveals that mink can spread the virus rapidly between animals and that the disease in mink is due to the viral infection and damage to tissues of the upper and lower respiratory system. The determination that mink are susceptible to SARS-CoV-2 indicates the need for strict biosecurity measures on mink farms to remediate mink-to-mink and human-to-mink transmission for the protection of mink, as well as prevent potential transmission from mink to humans.

## Introduction

Since December 2019, worldwide spread of a novel coronavirus designated as severe acute respiratory syndrome coronavirus 2 (SARS-CoV-2) has resulted in significant human disease, death, and economic loss [1]. Phylogenetic evidence suggests that SARS-CoV-2 may have jumped from the Intermediate Horseshoe Bat (*Rhinolopphus affinis*) to human beings, likely via an undetermined intermediate host [2–4]; if proven, this is an example of a generalist coronavirus broadening its host range. Other broadening coronavirus events in recent history include the 2002 emergence of Severe Acute Respiratory Syndrome – associated coronavirus (SARS-CoV) from a wildlife bat reservoir in China [5], and the 2012 emergence of Middle Eastern Respiratory Syndrome (MERS) from wild bats in Saudi Arabia [6]. SARS-CoV has a broad susceptible host range including naturally infected human beings, civet cats and raccoon dogs, and experimentally infected rhesus macaques, ferrets, mice, cats and hamsters [7–13]. Similarly, MERS is found in animal reservoir hosts such as bats, and dromedary camels [14]. As countries continue to modify infection control and public health strategies for containment of SARS-CoV-2, sources of viral transmission from domestic and wildlife animal reservoirs are of great interest. Natural and experimental SARS-CoV-2 infection studies demonstrate susceptibility of rhesus macaques, cats, dogs, ferrets, mice, tree-shrews, Egyptian fruit bats and Syrian guinea pigs and mink to the virus with variable permissiveness and expression of clinical disease; while pigs, poultry and cattle do not appear to be susceptible [15–25]. Investigations into the distribution of virus in experimental infected cats, ferrets and macaques demonstrate that viral RNA can be detected by polymerase chain reaction (PCR) in many organ systems, most notably the upper respiratory tract, lung and intestines [15,16,26]. SARS-CoV-2 viral proteins have been identified by immunohistochemistry (IHC) in nasal turbinates, trachea, lung and the lamina propria of intestines of experimentally infected ferrets [16] and lung, mediastinal lymph nodes and intestines of macaques [26]. Infectious viral particles have been recovered from nasal turbinates, nasal fluid, saliva, and lungs, but not from trachea, kidney and intestinal tissues of experimentally infected ferrets [16].

While these experimental studies provide valuable information regarding disease pathogenesis and possible animal reservoirs, the true risk of SARS-CoV-2 transmission between these species and human beings in natural settings remains undetermined. Natural SARS-CoV-2 infections have been reported in domestic dogs and cats, as well as large cats and great apes in zoological facilities [27–31]. The origin of infections in these settings has been attributed to human SARS-CoV-2 transmitting to animals. SARS-CoV-2 infections have also been reported in farmed mink worldwide, including the United States [32–36]. The index report from the Netherlands indicated respiratory disease and increased mortality in two mink farms in the Netherlands due to natural SARS-CoV-2 infection [17]. The Netherland’s mink outbreak investigation revealed viral RNA and protein present in multiple organ systems, most consistently detectable in respiratory system [17,37], and rapid transmission between mink in the mink facilities. Full-length viral genome sequencing from farmed mink SARS-CoV-2 outbreaks in the Netherlands and Denmark suggested novel virus variants with transmission between mink and humans and potential increased possibility of spread in this environment [34,38]. A new SARS-CoV-2 strain called “Cluster 5”, was identified in mink in Denmark that was also present in the human population raising concerns of a higher risk of people working on mink farms.

In August 2020 multiple mink farms in Utah experienced a sudden increase in animal mortality attributed to natural SARS-CoV-2 infection. The epidemiological information associated with the outbreak, gross and histopathologic lesions, tissue distribution of viral RNA, and genomic sequencing of the virus are described herein. The report details a large-scale natural infection outbreak of SARS-CoV-2 resulting in significant inter-animal transmission, disease and death in a susceptible animal species in the United States. Subsequent to this outbreak there have been reports of SARS-CoV-2 infection on mink farms in Oregon, Wisconsin and Michigan [39].

## Results

### Premise and animal information

The outbreak of disease associated with SARS-CoV-2 infection began in August 2020. Five mink farms were included in this investigation in which two farms had a common producer and the others were operated independently. Three premises had common labor between farms and were in close physical proximity (approximately 400 meters). Farms each had perimeter fences, locked gates, and access only to authorized personnel. Mink were housed in roof-covered sheds with ventilation to the outdoors through open side walls. Adult animals were held either individually or coupled in wire mesh cages with an approximately 1-inch space between cages to prevent inter-animal aggression, but nose-to-nose contact between neighboring animals was possible. Diets were comprised of offal and a carbohydrate source and were mixed in two distinct kitchens distributed daily to the five farms. Watering systems were variable between premises and animals had either individual nipple waterers, individual water dishes or a trough system. Mink were vaccinated annually against *Clostridium botulinum*, mink enteritis virus, canine distemper virus, and *Pseudomonas aeruginosa*. Aleutian mink disease virus was intermittently identified as a cause of disease on the farms and considered a possible comorbidity. Wildlife, including skunks and raccoons, were intermittently observed on the premises and eliminated on an as-needed basis. Feral cats were commonly present on premises to assist with rodent control.

Clinical disease, including death, was observed in adult breeding animals ranging in age from 1-5 years, while young-of-the-year kits were overwhelmingly unaffected by the virus. The first sign noted by producers was an abrupt increase in the overall mortality rate. The mortality rate ranged from 35-55% in the adult-aged mink, which normally ranges between 2 and 6%. On one premise the mortality rate in female mink was 1.8 times greater than males. An increase in respiratory effort was notable in diseased mink characterized by gasping or increased abdominal effort. Upper respiratory signs included nasal and ocular discharge (Fig 1a), and coughing was present, but was variable between farms. There was no report of gastroenteritis. Survival with resolution of respiratory disease was observed in some animals without observable lasting effects, however the frequency of this occurrence is unknown. The source of the virus was presumed to be due to reverse zoonosis of the virus from infected workers [31].

**Figure 1:**
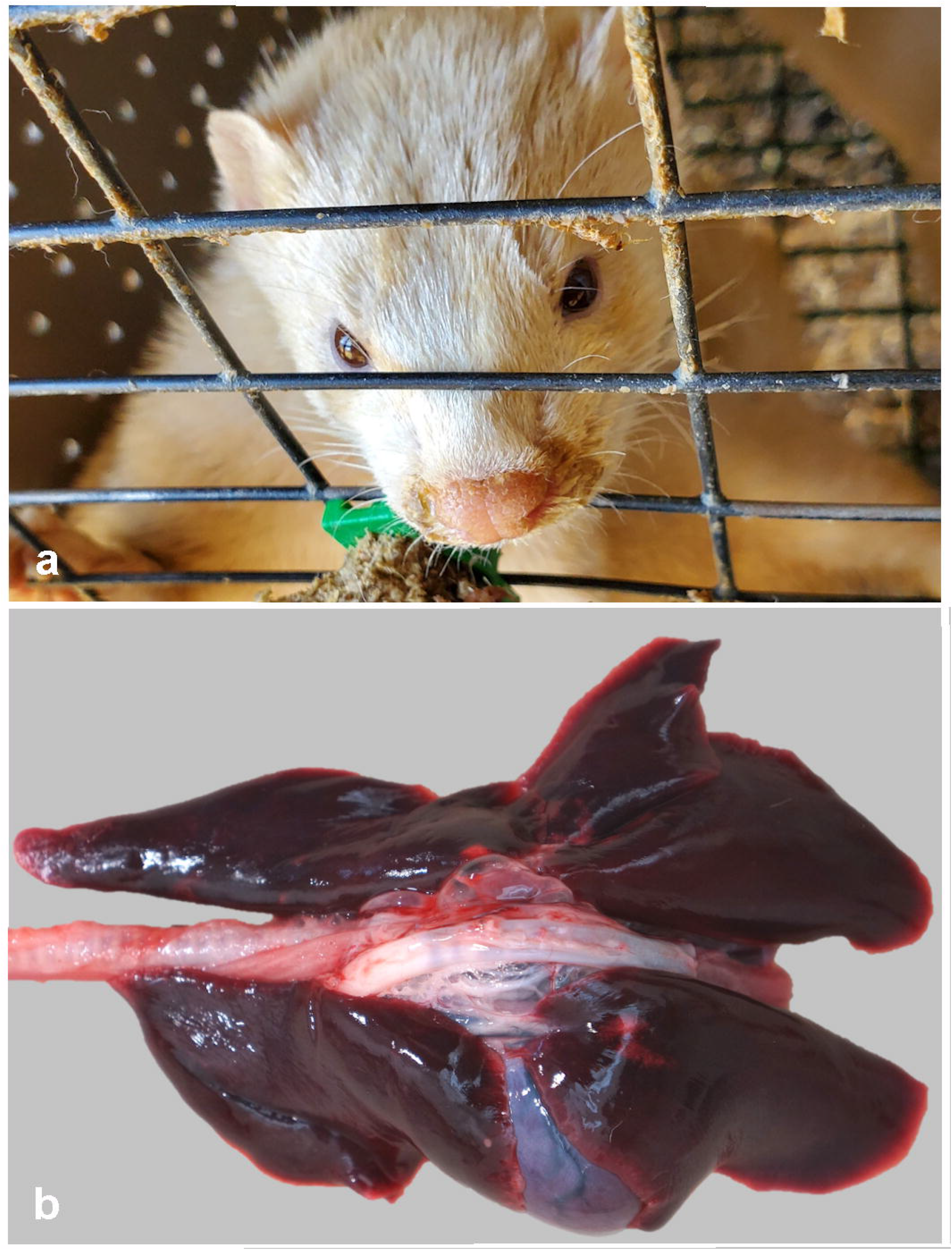
Clinical and gross necropsy findings in SARS-CoV-2 infected mink. a. A mucopurulent nasal discharge, indicative of rhinitis, stains the fur surrounding the nares in a SARS-CoV-2 infected mink. b. Gross image of severe pulmonary congestion and edema of an infected mink.

### Pathology

A total of 20 mink, both female and male, from five farms were necropsied (examined postmortem). Given the clinical suspicion of SARS-CoV-2 infection and the potential risk to human prosectors, necropsies were performed with personal protective equipment in accordance with biosafety level 3 practices (conducted in Class II biosafety cabinet with appropriate primary barriers and personal protective equipment including clothing, gloves, eye, face and respiratory protection). Mink were generally in good body condition based on adipose tissue stores and muscling. In all mink, lung lobes were uniformly (most common) or variably dark red, heavy, and failed to collapse (Fig 1b). Abundant clear fluid escaped when lung lobes were incised, and tracheas contained variable amounts of white froth (pulmonary edema).

Histopathological examination revealed multifocal interstitial and perivascular pneumonia (Fig 2a) and variable amounts of alveolar edema (Fig 2b) in all mink. Occasional fibrin strands overlay necrotic alveolar pneumocytes. Proliferative type II pneumocytes infrequently lined other alveolar septa (Fig 2c). Low to moderate numbers of neutrophils and macrophages plus moderate amounts of fibrin were in multiple alveolar spaces (Fig 2d). In nearly all pulmonary arterioles, edema fluid and moderate numbers of lymphocytes and plasma cells widely separated collagen fibers of the tunica adventitia. Sporadic vessels had mural fibrinoid degeneration. Additional findings included mild, diffuse, catarrhal to necrotizing enterocolitis (5/20 mink), moderate, multifocal, splenic lymphoid necrosis (5/20 mink), severe acute centrilobular hepatic congestion (4/20 mink), focal perivascular lymphocytic meningitis (1/20 mink), severe necrotizing and suppurative bridging centrilobular hepatitis (1/20 mink) and myocardial interstitial fibrosis and fatty infiltration (1/20 mink). Severe suppurative rhinitis with multifocal attenuation and loss of epithelial cells was noted on histopathological examination of nasal turbinates from two mink.

**Figure 2:**
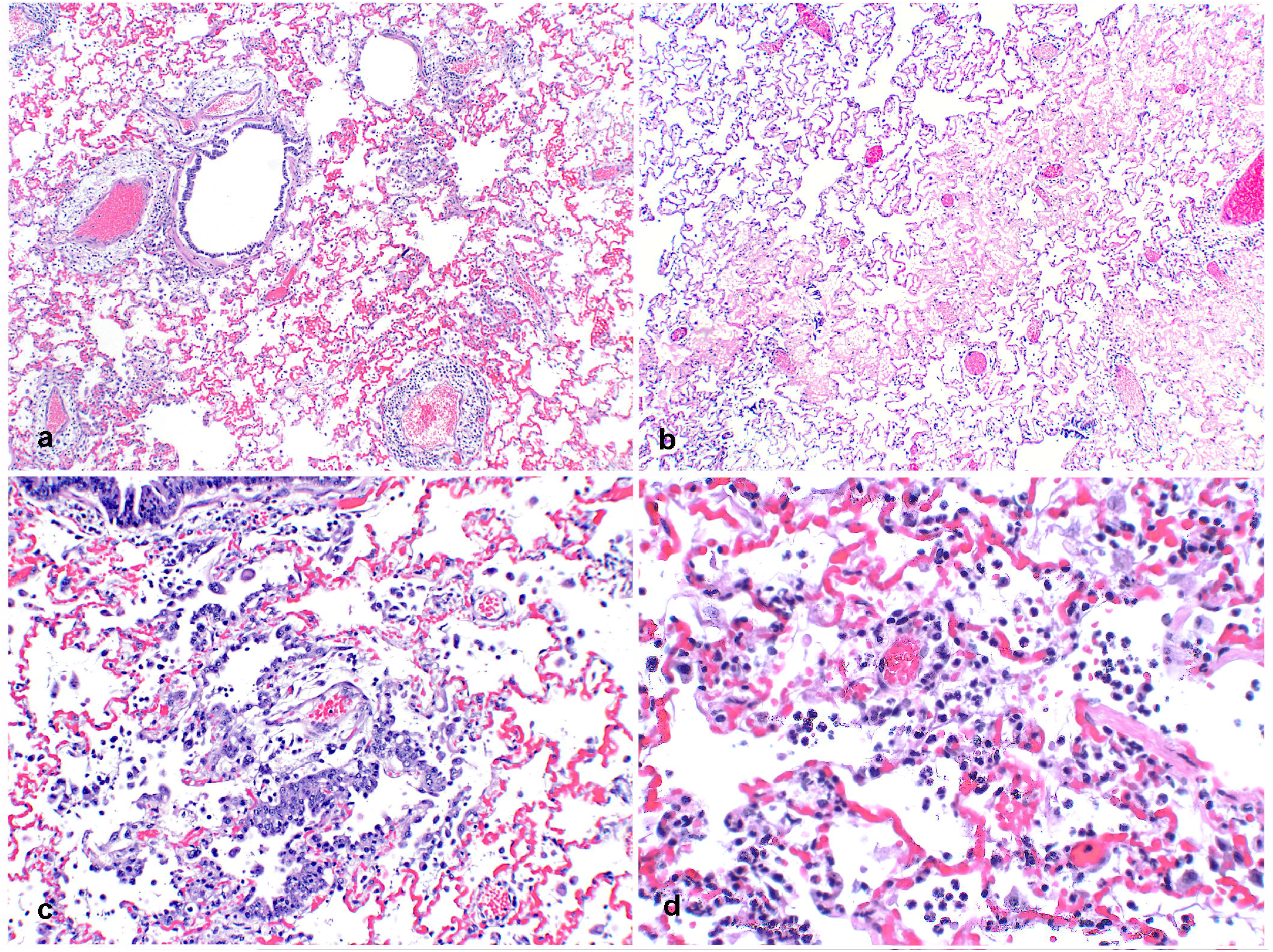
Pulmonary histopathology of SARS-CoV-2 infected mink. a. Lung from an adult mink with large cuffs of mononuclear inflammatory cells and edema multifocally surrounding pulmonary vessels. 20x H&E. b. Alveolar spaces are multifocally filled with eosinophilic edema fluid. 40x H&E. c. Bronchioles are lined with proliferative, slightly disorganized hyperplastic epithelium and type II pneumocyte hyperplasia is present in alveoli associated with increased intra-alveolar inflammation. 100x H&E. d. Neutrophils, fewer macrophages, and strands of fibrin are multifocally present in alveoli. 200x H&E.

### Tissue Distribution of SARS-CoV-2 by RT-PCR

The initial detection of SARS-CoV-2 infection was from deep nasopharyngeal swabs and fresh lung tissue from five necropsied mink from two farms by RT-PCR. Subsequent to this initial diagnosis multiple additional tissues from four necropsied animals were collected in Trizol for further investigation of viral tissue distribution by RT-PCR (designated mink 1-4). Viral RNA was detected in many tissues from multiple mink (Fig. 3a). Tissues where SARS-CoV-2 viral RNA was consistently detected between animals included nasal turbinates and lung, where nasal turbinates had a lower cycle threshold detectability than lung. Other tissues where viral RNA was detected included the retropharyngeal lymph node (3/4 mink), tracheobronchial lymph node (3/4 mink), squamous tissue from the distal nose (3/4 mink), paw pads (3/4 mink), and brain (3/4 mink). Detectible viral RNA was observed in other tissues with less frequency between animals. Once it was identified that nasal turbinates from two of the initially sampled mink (mink 3 and 4) had very low Ct detectability by RT-PCR (interpreted as a high viral load), RT-PCR was performed on formalin-fixed paraffin embedded (FFPE) sections of nasal turbinates from two additional mink (mink 5 and 6), where SARS-CoV-2 RNA was also detected.

**Figure 3:**
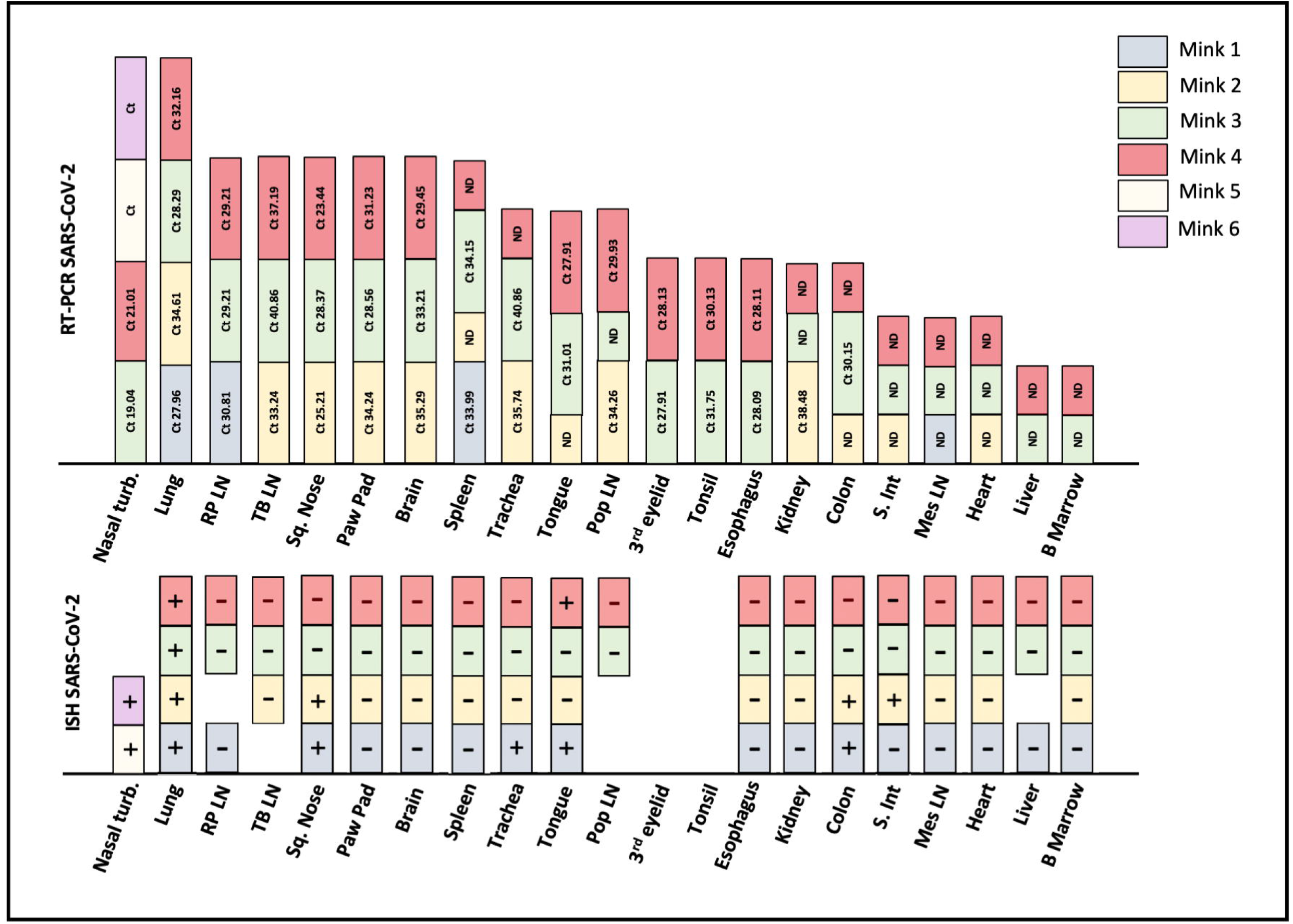
Detection of SARS-CoV-2 RNA in tissues by RT-PCR and ISH. a. Tissues where viral RNA was not detected are represented by “ND” and tissues not collected/not tested are represented by an empty space. CT, cycle threshold. b. Tissues in which viral RNA was detected by a chromogenic signal by ISH are represented with a “+”, and “-” when not detected.

### Virus Sequence Analysis

Whole genome viral sequences from all of the mink farms were identical. Mutational analysis was performed using the GISAID EpiFlu^™^ Database CoVsurver: Mutation Analysis of hCoV-19 at https://www.gisaid.org/epiflu-applications/covsurver-mutations-app. The SARS-CoV-2 viral sequences from the mink were in GISAID clade GH, with mutations at T85I-NSP2, S1205L-NSP3, G37E-NSP9, P323L-NSP12, T91M-NSP15, D614G-spike, N501T-spike, Q57H-NS3, H182Y-NS3, A38S-M, T205I-N, and Q289H-N as compared to hCoV-19/Wuhan/WIV04/2019. See Table 1 for all SNPs and aa mutations.

**Table 1:**
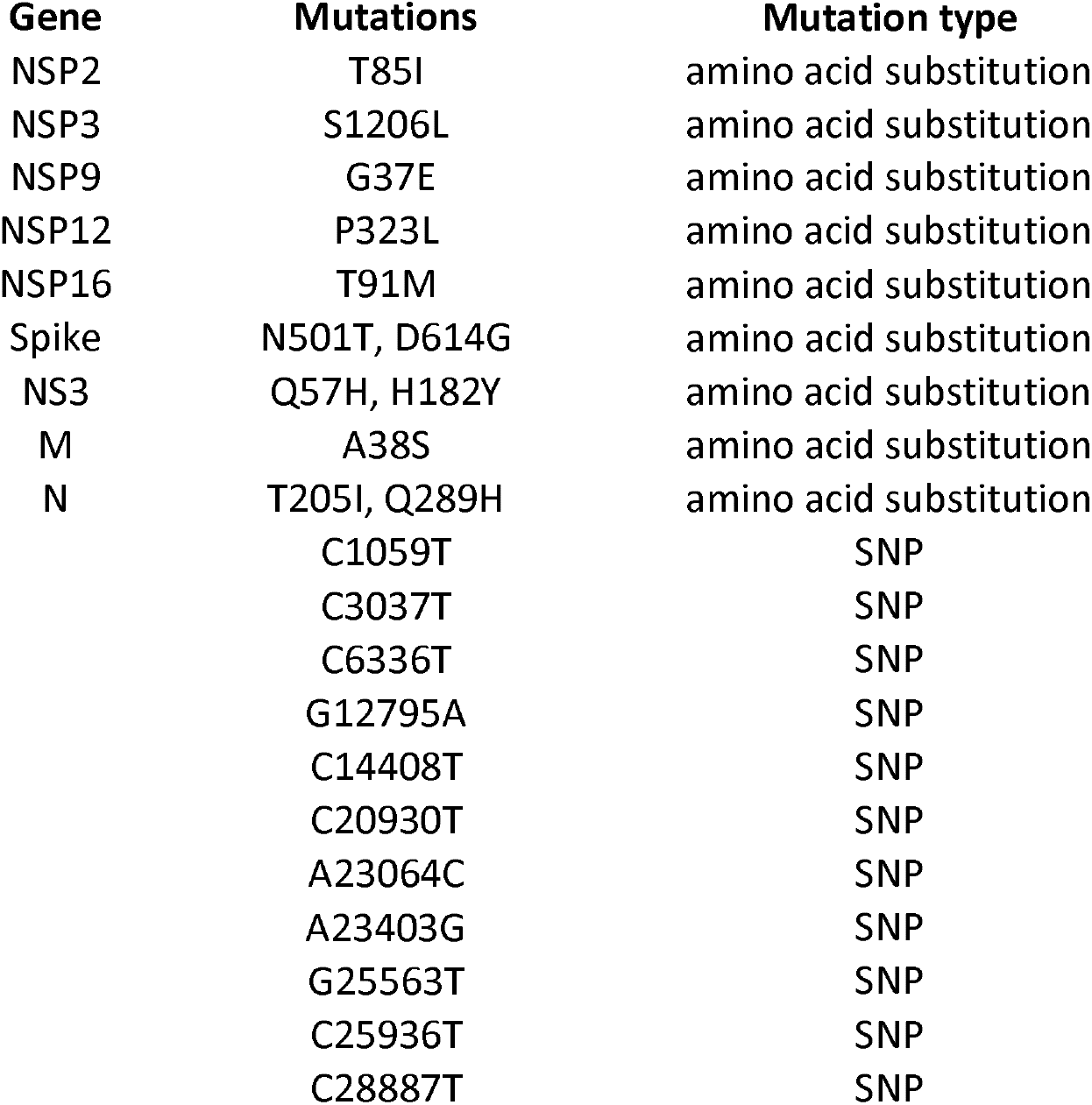
Mutations identified in the genome of Utah mink SARS-CoV-2 isolates. SNPs and nonsynonymous mutations identified. Amino acid and codon numbering is relative to Wuhan-Hu-1.

### Cellular Distribution of viral RNA in tissues

SARS-CoV-2 RNA was detected by chromogenic *in situ* hybridization in multiple FFPE tissues (Fig 3b). The nasal turbinates and nasal passages of the two mink in which tissues were available for evaluation had abundant positive staining for viral RNA in the suppurative and catarrhal exudate within the nasal passages (Fig 4a-c), as well as in the respiratory epithelial cells of the most caudal nasal passage overlying nasal mucous glands (Figs 4d-f). In 4/4 mink there was positive detection of viral RNA in pulmonary bronchial epithelial cells or multifocally within the interstitium (Figs 4g-h). There was also positive detection of viral RNA in the tracheal epithelial cells of one mink 1 (Fig 4i-j). Other tissues where viral RNA was observed included the most superficial surface of the distal squamous nose (2/4 mink), the surface of the tongue (1/4 mink), and very little detection in the lumen of the colon (2/4 mink) and small intestine (1/4 mink). Thryoid gland, adrenal gland, eye, ovary, uterus and pancreas were examined from 1 mink by ISH (data not included in Fig 3b) in which viral RNA was not detected. Positive and negative tissue and reagent controls, as described in Materials and Methods, performed as expected.

**Figure 4:**
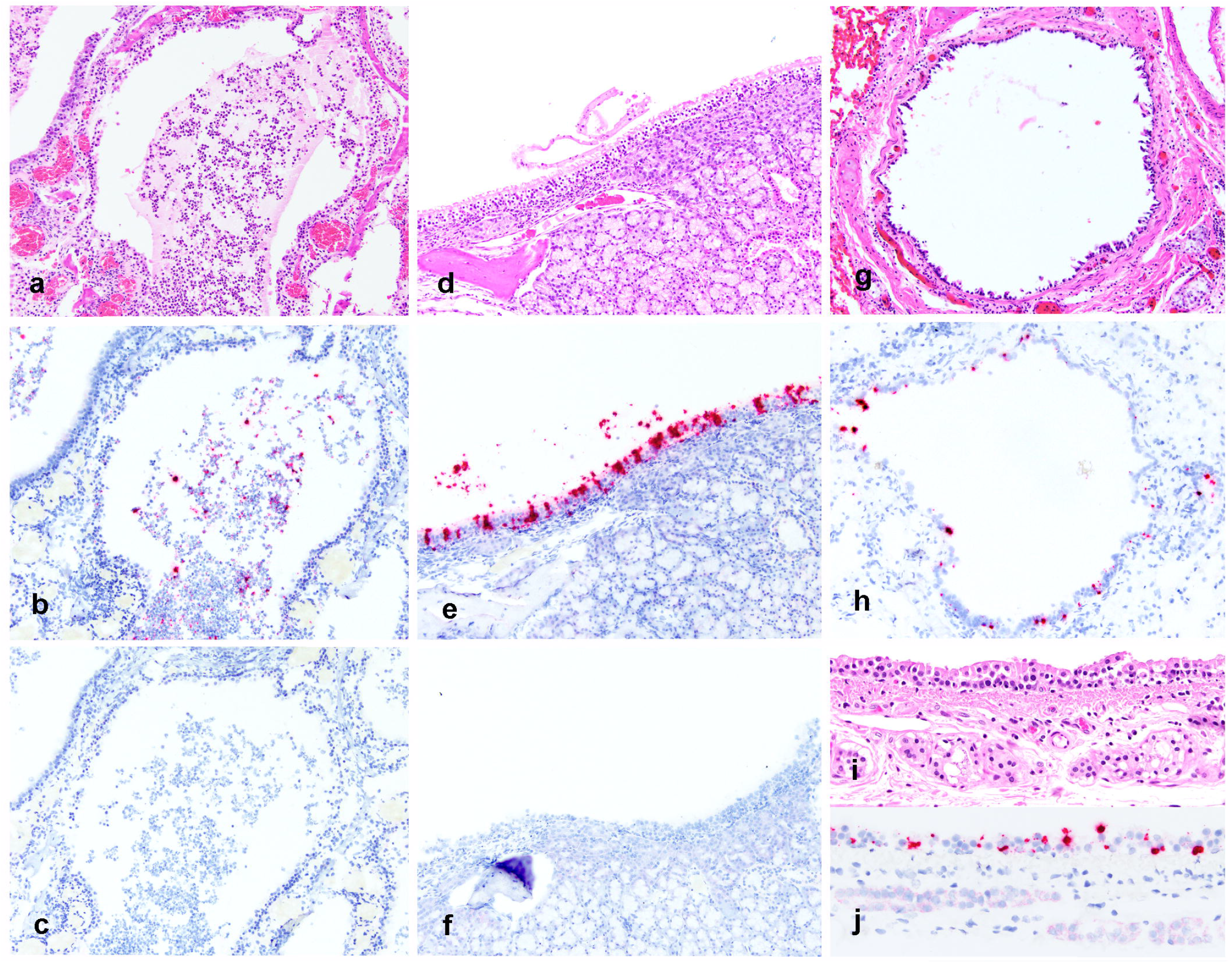
Detection of SARS-CoV-2 RNA in tissues by ISH. Figures a-c. Nasal turbinate samples from SARS-CoV-2 infected mink 5. a. H&E image of suppurative and histiocytic rhinitis filling the nasal passage. b. Detection of SARS-CoV-2 RNA in the nasal exudate. c. No detection of FIPV RNA in nasal turbinate of SARS-CoV-2 infected mink (negative control). Figures d-e d. Nasal turbinate samples from SARS-CoV-2 infected mink 5 demonstrating mild rhinitis in the caudal nasal passage and mild disorganization of respiratory epithelial cells e. Detection of SARS-CoV-2 viral RNA within respiratory epithelial cells of mink 5. f. No detection of FIPV RNA in nasal turbinate epithelial cells of SARS-CoV-2 infected mink (negative control). g-h Lung samples from SARS-CoV-2 infected mink 1. g. H&E image of bronchus with attenuation and multifocal loss of respiratory epithelial cells. h. Detection of SARS-CoV-2 viral RNA within respiratory epithelial cells of the bronchus. i-j Trachea samples from SARS-CoV-2 infected mink 1. i. H&E image of trachea with mild attenuation and multifocal disorganization of respiratory epithelial cells. j. Detection of SARS-CoV-2 viral RNA within respiratory epithelial cells of the trachea.

### Discussion

In this report we show that mink are highly susceptible to SARS-CoV-2 infection with high mortality in a natural farm production setting. Furthermore, we describe the pathology and tissue distribution of the virus in infected animals. Since the emergence of SARS-CoV-2, mink have been the only animal species identified to develop significant disease and mortality associated with infection in the United States. The findings were associated with reverse zoonotic transmission (from humans to mink) similar to other SARS-CoV-2 infections reported in animals. The abundant mink-to-mink transmission occurring on multiple farms with high morbidity and mortality highlighted concerns regarding propagation of viral mutants with greater fitness and virulence. In a recent Denmark investigation, SARS-CoV-2 infected mink, many that were asymptomatic, were suggested to serve as transmission vectors of a new mutated strain of virus to humans [34]. Preliminary viral molecular phylogeny and epidemiology studies of the Utah farms described herein has not identified the SARS-CoV-2 mutations associated with mink-to-human transmission in the Danish study.

The outbreaks of SARS-CoV-2 in farmed mink in April 2020 in the Netherlands and Denmark showed robust viral transmission, similar to what we have described here [17,34,37],. Mortality rates in our Utah outbreak were much higher (up to 55%), compared to the Netherlands outbreak, which reported 2.4% mortality at greatest, and the Danish outbreak, which showed minimal clinical disease and mortality [33,34]. Such a substantial difference in mortality may be due to the population of mink considered in the mortality rate (only adults were considered in this case, while young animals may be have included in the Netherland report), or other reasons such as differences in housing and management, comorbidities (such as infection with Aleutian Disease virus), or viral virulence. Since the outbreak of SARS-CoV-2 in the Utah mink there have been infections in multiple other mink farms in the United States including Oregon, Wisconsin and Michigan [39].

In our pathology investigations the most significant findings were observed in the respiratory tract, and death was attributed to pulmonary failure and edema. Histologically, the respiratory changes were typical of viral interstitial pneumonia with alveolar damage, consistent with the pulmonary histopathology described in the Netherlands mink outbreak [37]. One interesting histopathologic finding of note in our case not described in the Netherland outbreaks was the presence of perivascular mononuclear inflammatory cells, edema and rare vascular wall fibrinoid necrosis (vasculitis), which has been described in humans and experimentally infected ferrets [15,40–42]. In a recent report of describing the pulmonary pathology from human Covid-19 deaths, a key histologic feature of was the presence of increased numbers of perivascular T-lymphocytes (termed pulmonary vascular endothelialitis), though this feature did not definitively distinguish it from influenza pneumonia [42]. Given some of the striking similarities between the pulmonary histopathology of SARS-CoV-2 infected mink and humans, and abundance of virus in the upper respiratory tract between species, mink should be considered as a very good natural disease model of human Covid-19 disease.

The finding of severe suppurative and catarrhal rhinitis observed in the infected Utah mink was also an interesting finding. Rhinitis has been described in association with SARS-CoV-2 infection in experimentally infected cats, but the nature of the inflammation was described as mononuclear rather than suppurative [15]. Examination of the nasal conchae in the Netherlands mink report revealed swelling and degeneration of epithelial cells with diffuse loss of cilia and mild inflammation, which wasn’t further characterized [37]. In any case, significant differences were observed in the nasal inflammation between these two outbreaks, which should be addressed in future investigations.

The tissue distribution of virus investigated by RT-PCR reported herein revealed fairly consistent detection of viral RNA in upper and lower respiratory tissues. Interestingly, RT-PCR also detected viral RNA in the brain, spleen, and various lymph nodes of multiple mink. By ISH, viral RNA was localized to respiratory epithelial cells of nasal turbinates, trachea and bronchi with multifocal detection in the pulmonary interstitium of some mink. These cellular localization findings are similar to the Netherlands investigation, which demonstrated viral antigen in epithelial cells in the same locations [37]. In experimentally infected cats, ferrets and Syrian hamsters the distribution of viral antigen localization was similar [15,21]. In this case we also identified viral RNA present on superficial epithelial surfaces of the distal nose, tongue and rarely within the lumen of the intestines by ISH, which we interpret as likely shedding from the infected nasal passage and passive surface accumulation or ingestion. This finding is interesting and may suggest that infectious virions are present on superficial epithelial surfaces and are potential sources of viral transmission. Detection of intact infectious virions would be necessary to prove this hypothesis. There were discrepancies in the tissue distribution of viral RNA as detected by RT-PCR and ISH in this report, which warrants further investigation. These differences could be due to a greater detection sensitivity by RT-PCR, contamination of samples during collection at necropsy and detection by RT-PCR, or viremia with rare or inconsistent detection in various tissue systems. In a recent study investigating the utility of RNA-ISH, immunohistochemistry and RT-PCR in humans infected with SARS-CoV-2, ISH had a sensitivity and specificity of 86.7% and 100% respectively compared to RT-PCR [43]. Additionally, they report that RT-PCR and ISH consistently demonstrated the presence of viral RNA within pulmonary tissues, where viral RNA was not detected in any extrapulmonary tissues by either method. Another likely contributor to the differences we report here may be due to the small sample size of mink investigated, which is considered a limitation of this report.

Omitting 42 ambiguous bp reads in the stable NSP-9 region, all five viral whole genome sequences from the mink isolates were 100% identical with three human SARS-CoV-2 GenBank accessions from Washington State, MW474211, MW474212, and MW474111, and mutations discovered via GISAID analysis are identical between the mink isolates and these three human isolates. GISAID differentiates COVID-19 into three major clades: Clade S, Clade V and Clade G (originally prevalent in North America, Asia/Europe, and Europe, respectively), based on NS mutations at NS8_L84S, NS3_G251V and S_D614G, respectively [44]. The G clade was subsequently divided into GR clade containing N_203-204: RG>KR and GH clade with NS3_Q57H aa substitutions [45]. The Utah mink isolates fall into clade GH. Analysis via the CoV-GLUE website at http://cov-glue.cvr.gla.ac.uk/#/home {CoV-GLUE: A Web Application for Tracking SARS-CoV-2 Genomic Variation Joshua B Singer, Robert J Gifford, Matthew Cotten and David L Robertson Preprints 2020, 2020060225 https://doi.org/10.20944/preprints202006.0225.v1}classifies this virus in the B.1 lineage of the Rambaut et al. lineage system. {Andrew Rambaut, Edward C Holmes, Áine O’Toole, Verity Hill, John T McCrone, Christopher Ruis, Louis du Plessis and Oliver G Pybus Nature Microbiology 2020 https://doi.org/10.1038/s41564-020-0770-5}. This lineage originally comprised the Italian outbreak before spreading to Europe and other parts of the world.

Twelve nonsynonymous sequence mutations were identified in the SARS-CoV-2 genome from the Utah mink isolates. The polyprotein ORF1ab T85I-NPS2 mutation is most common in the USA (56% of phase 2 viruses) and has spread to at least 37 countries during phase 2 of the pandemic [46]. The P323L-NSP12 mutation in the viral polymerase gene coevolved with the D614G-spike mutation also present in this mink strain to become the most prevalent variant in the world. The G614 variant of the spike is more infectious than the original Wuhan D614 variant. Success of the P323L/ G614 variant suggests that the P323L mutation adds to the virulence of the G614 spike variant, although without increasing patient mortality. [47]. In addition to the highly prevalent D614G mutation, the mink isolate had a rare N501T spike mutation. GISAID reports that this mutation is related to host change and antigenic drift. N501T is located in the Receptor Binding Domain (RBD) of the spike glycoprotein, resulting in a moderate increase in ACE2 binding [48,49]. The NS3_Q57H mutation is common in the USA and is predicted to be deleterious [50]. S1205L-NSP3, T91M-NSP15, H182Y-NS3, Q289H-N, and A38S-M are rare mutations of unknown significance. Utah mink did not share other spike RBD mutations Y453F and F486L found in mink, nor did they have any of the common mutations reported from other mink throughout the world. These included five nsp2 aa substitutions (E352Q, A372V, R398C, A405T, and E743V), four in the nsp3 papain-like proteinase domain (P1096L, H1113Y, I1508V, and M1588K) one in the nsp5 3C-like proteinase domain (I3522V), one in the nsp9 RNA/ DNA binding domain (G4177E or R) one in the nsp15 poly(U) specific endoribonuclease domain (A6544T), two in the nsp12 RNA-dependent RNA polymerase domain (M4588I and T5195I), and two in the nsp13 helicase domain (I5582V and A5770D). {Elaswad A, Fawzy M, Basiouni S, Shehata AA. Mutational spectra of SARS-CoV-2 isolated from animals. PeerJ. 2020 Dec 18;8:e10609. doi: 10.7717/peerj.10609. PMID: 33384909; PMCID: PMC7751428.} The more uncommon spike RBD N501T mutation from Utah mink has been found in four emergences within three lineages of mink samples. {Recurrent mutations in SARS-CoV-2 genomes isolated from mink point to rapid host-adaptation Lucy van Dorp, Cedric CS Tan, Su Datt Lam, Damien Richard, Christopher Owen, Dorothea Berchtold, Christine Orengo, François Balloux bioRxiv 2020.11.16.384743; doi: https://doi.org/10.1101/2020.11.16.384743}

With the exception of the common D614G mutation, the Utah mink have none of the multiple spike protein changes (deletion 69-70, deletion 145, N501Y, A570D, D614G, P681H, T716I, S982A, D1118H) defining human UK variant VUI 202012/01, which may have increased transmissibility compared to other variants, nor does it have any of the mutations defining novel human South African variant 501Y.V2 (spike RBD K417N, E484K, and N501Y)

In conclusion, our results indicate that mink are susceptible to SARS-CoV-2 infection and can readily transmit the virus between animals. Infected animals suffer from severe respiratory disease, similar to that which has been described in humans, as well as other experimentally infected animals. Further investigations should focus on investigating the immunology and vascular pathology associated with the development of disease in mink to potentially extrapolate findings for human health and other animals. The Utah mink SARS-CoV-2 strain is unique among mink and other animal strains sequenced to date. Identical strains found in Washington state humans may reflect zooanthroponosis, and to date there is no evidence that viruses adapted to mink will impact human SARS-CoV-2 evolution. However, monitoring of mutations located within the RBD of the SARS-CoV-2 spike protein in mink is important for studying viral evolution and host-adaptation. Between August 2020 and the end of January 2021 the N501T mutation increased in frequency of sequenced isolates in the United States from .01% to .30%, similar to the increase in N501Y mutations. Lastly, strict biosafety measures are warranted on mink farms to decrease viral transmission between animals and risk of transmission to humans, as well as decreasing animal losses due to SARS-CoV-2 infection.

## Materials and Methods

### Pathology

Deceased mink were submitted to the Utah Veterinary Diagnostic Laboratory for investigation of the cause of death. In most cases animals died acutely due to natural infection, and less commonly were euthanized by cervical dislocation when humane euthanasia was warranted according to Fur Commission USA standards. The mink were housed as distinct separate, private operations that fall outside of the IACUC approval required at universities. All farms were members of the Utah Fur Breeders Association, which is under the Fur Commission USA and all members follow standard guidelines for the operation of mink farms in the United States, which includes best practices for care, biosecurity and euthanasia.

At necropsy all body systems were examined by an ACVP-board certified anatomic pathologist (TB) and one anatomic pathology resident (MC) at the Utah Veterinary Diagnostic Laboratory.

A full complement of tissues were collected from twenty mink and fixed in 10% neutral buffered formalin. Formalin fixed tissues were dehydrated in ethanol, embedded in paraffin wax, sectioned at 4 μm, and stained with hematoxylin and eosin using standard histochemical techniques. For SARS-CoV-2 RT-PCR, an extended list of tissues were collected and placed into TRIZOL Reagent (ThermoFisher Scientific, Waltham, MA); Oronasal swabs were placed in viral transport medium (PrimeStore MTM; LongHorn Diagnostics).

### Swab sample extraction method

Total nucleic acid was extracted from samples in 1 mL of PrimeStore MTM [LongHorn Diagnostics] using MagMAX^™^-96 Viral RNA Isolation Kit, per the manufacturer’s instructions.

### Tissue sample extraction method

RNA was extracted from fresh tissue samples in TRIZOL Reagent and formalin fixed paraffin embedded tissues using TRIzol^™^ reagent [ThermoFisher, Waltham, MA 02451], per the manufacturer’s instructions. FFPE tissues were cut in 10um sections and heated at 65°C for 10 minutes in Trizol prior to RNA extraction using MagMAX^™^-96 Viral RNA Isolation Kit, per the manufacturer’s instructions.

### RT-PCR conditions

Reverse transcriptase (RT) real-time PCR to the SARS-CoV-2 RNA-dependent RNA polymerase gene (RDRp) was performed as previously described using primers SARS-CoV-2 primers RdRp_SARSr-F2 5’-GTGARATGGTCATGTGTGGCGG-3’ and COVID-410R 5’-CCAACATTTTGCTTCAGACATAAAAAC-3’ [51], using TaqMan Fast Virus 1-Step Master Mix Kit [Thermo Fisher]. RNA amplification was done using ABI 7500 Fast (ThermoFisher, Waltham, MA 02451). Controls included positive extraction control (RdRp_GATTAGCTAATGAGTGTGCTCAAGTATTGAGTGAAATGGTCATGTGTGGCGGTTCACTATATGTTAAACCAGGTGGAACCTCATCAGGAGATGCCACAACTGCTTATGCTAATAGTGTTTTTAACATTTGTCAAGCTGTCACGGCCAATGTTAATGCACTTTTATCTACTGATGGTAACAAAATTGCCGATAAGTATGTCCGCAATTTAC, negative extraction control (PCR water), positive amplification control (SARS-CoV-2 whole genome RNA), and negative amplification control (No template control). Graphs and tabular Ct results were reviewed on the ABI 7500 program. Unknown samples were considered positive if they rose above the threshold by cycle 45. All others were considered negative.

### Whole Genome Sequencing

Libraries for the whole genome sequencing were generated using the Ion AmpliSeq Kit for Chef DL8 and Ion AmpliSeq SARS-CoV-2 Research Panel (Thermo Scientific, Waltham, MA). Libraries were sequenced using an Ion 520 chip on the Ion S5 system using the Ion 510^™^ & Ion 520^™^ & Ion 530^™^ Kit. Sequences were assembled using IRMA v. 0.6.7 and visually verified using DNAStar SeqMan NGen v. 14. Mutational analysis was performed using the GISAID EpiFlu^™^ Database CoVsurver: Mutation Analysis of hCoV-19 at https://www.gisaid.org/epiflu-applications/covsurver-mutations-app.

### Visualization of genomic material in tissues

*In situ* hybridization utilized RNAscope (Advanced Cell Diagnostics, Hayward, CA) technology to visualize the presence and location viral RNA in tissues harvested from infected mink. A set of anti-sense SARS-CoV-2 specific RNA probes comprised of 20 Z pairs targeting nucleotides 21,631-23,303 of the spike viral glycoprotein gene (Genbank accession number NC_045512.2) was developed by Advanced Cell Diagnostics (ACD) and performed as previously described [52]. This assay was performed according to manufacturer’s protocols for RNAscope 2.5 HD Red Detection Kit (ACD) with the following specific conditions. Fresh tissues from four SARS-CoV-2-positive and two SARS-CoV-2 negative mink were fixed in 10% buffered formalin, embedded in paraffin wax, and sectioned at 4um on positively charged glass slides. Samples were slowly submerged in lightly boiling Target Retrieval Solution (ACD) for 15 minutes, followed by application and incubation of Protease Plus (ACD) at 40°C for 20 minutes. In addition, two SARS-CoV-2 negative mink were selected from the UVDL tissue achieves as negative controls. A probe specific for a feline infectious peritonitis virus (FIPV) RNA also generated by ACD as positive and negative controls. FFPE tissues from a domestic cat with peritonitis due to FIPV-infection was used as positive assay control. Additionally, these FIPV-infected tissues, FFPE intestinal tissue from a bovine calf infected with bovine corona virus (confirmed by PCR), a coronavirus positive calf trachea and nasal turbinates from a domestic cat all were utilized as negative tissue controls and stained with the SARS-CoV-2 probe to investigate any cross-reactivity to these other coronaviruses and non-specific reactivity.

There was detection of viral RNA in inflamed splenic tissue from an FIPV infected cat, which served as an assay control. No viral RNA was detected (no cross-reactivity) in the negative control slides which included applying the SARS-CoV-2 probe to tissues (lung, lymph node, small intestine and colon) from one healthy adult mink that died of crush injuries prior to the emergence of SARS-CoV-2, spleen from an FIPV-infected cat, intestines and trachea from a bovine calf infected with bovine coronavirus, and nasal turbinates from a cat with suppurative rhinitis collect prior to the emergence of SARS-CoV-2. Additionally, there was no FIPV detection when this probe was applied to the SARS-CoV-2 positive mink nasal turbinates and lungs (Fig 4c,f).

## Acknowledgements

We would like to acknowledge the Utah mink breeders whose cooperation allowed this work. Additionally, we would like to acknowledge both the United States Food and Drug Administration Veterinary Laboratory Investigation and Response Network (VetLIRN) and the United States Department of Agriculture National Animal Health Laboratory Network (NAHLN) for their assistance in funding this project.

